# Divisome core complex in bacterial cell division revealed by cryo-EM

**DOI:** 10.1101/2022.11.21.517367

**Authors:** Lisa Käshammer, Fusinita van den Ent, Magnus Jeffery, Nicolas L. Jean, Victoria L. Hale, Jan Löwe

**Author notes:** Corresponding author: Jan Löwe. authors contributed equally (Lisa Käshammer, Fusinita van den Ent).

## Abstract

Cell division, or cytokinesis is a fundamental process of life and, in most bacteria, is driven by peptidoglycan synthesis at the septum^1^. It is catalysed by the divisome, a multi-protein complex with more than 20 components that spans the cell envelope in bacteria harbouring a cell wall^2^. Central to the divisome is the peptidoglycan-synthesising protein complex FtsWI, with the transglycosylase (TG) FtsW polymerising glycan strands from its substrate Lipid II^3,4^, and the transpeptidase (TP) FtsI crosslinking peptide stems, thus forming a covalent mesh between glycan strands^5,6^ (Fig. 1a). Septal peptidoglycan synthesis occurs after activation of the divisome glycosyltransferase-transpeptidase pair FtsWI^3^, in particular through an interaction with the heterotrimer FtsQBL^7^.

Here, we present the cryo-EM structure of the catalytic divisome core complex FtsWIQBL from *Pseudomonas aeruginosa* at 3.7 Å resolution. The structure reveals the intricate details of the periplasmic interfaces within FtsWIQBL, including the positioning of FtsI by the coiled coil of FtsBL, as well as a transmembrane domain containing FtsWIBL but not FtsQ. With our structure we are able to provide molecular mechanisms of a multitude of known mutations that interfere with divisome activation and regulation. Finally, we reveal a large conformational switch between presumably inactive and active states of the FtsWI core enzymes.

Our work is foundational for further structural, biochemical and genetic studies elucidating the molecular mechanisms of bacterial cell division. Since the divisome peptidoglycan synthase is essential for cell division in most bacteria, and is absent in eukaryotic cells entirely, it is a key target of important antibiotics and antibiotic development^8^, and we suggest that our structure will help to accelerate these efforts.

## Cryo-EM structure of the core divisome complex FtsWIQBL

To solve the structure of the core divisome complex, we initially purified the *Escherichia coli Ec*FtsWIQBL complex expressed in insect cells (Fig. S1a). As the *Ec*FtsWIQBL sample was heterogeneous, we were unable to solve its structure and hence purified the *Pseudomonas aeruginosa Pa*FtsWIQBL complex expressed in *E. coli* (Fig. 1b and S1b). Both *Ec*FtsWIQBL and *Pa*FtsWIQBL possess comparable transglycosylase activity, while the putative active site mutant *Pa*FtsW^D275A^IQBL is inactive (Fig. 1c). This is in accordance with previous data where the *Pa*FtsW^D275A^IQBL mutation caused filamentation in *P. aeruginosa* cells when overexpressed and displayed reduced transglycosylase activity *in vitro*^*3*^. Having confirmed that the purified *Pa*FtsWIQBL complex produced peptidoglycan strands, we proceeded with single-particle averaging cryo-EM and determined the structure of *Pa*FtsWIQBL to a final overall resolution of 3.7 Å (Fig. 1d, e, S1c-e, Table S1).

**Fig. 1:**
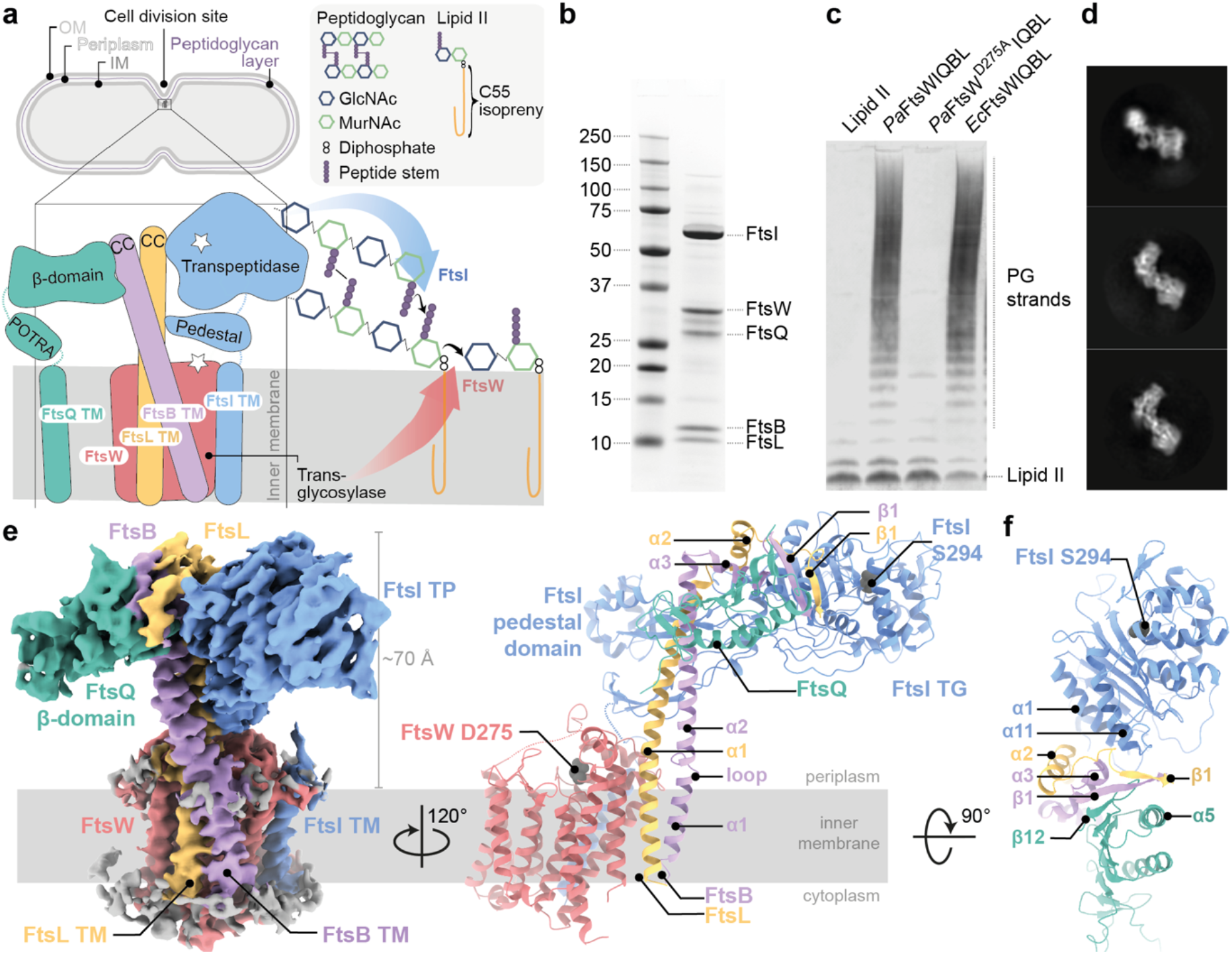
Biochemical and structural characterisation of the core divisome complex FtsWIQBL from *P. aeruginosa*. **a)** Septal peptidoglycan synthesis by FtsWIQBL during Gram-negative bacterial cell division. The transglycosylase FtsW (red), and transpeptidase FtsI (blue) bind the non-enzymatic subcomplex FtsQBL (green, violet and yellow, respectively). The complex contains 14 transmembrane helices – ten from FtsW and one each from FtsIQLB. The transglycosylase FtsW catalyses the polymerisation of GlcNAc-MurNAc disaccharides from Lipid II. The transpeptidase FtsI crosslinks the peptides from the nascent chain to adjacent peptides in the peptidoglycan layer between residues three and four. OM: outer membrane, IM: inner membrane, GlcNAc: N-acetylglucosamine, MurNAc: N-acetylmuramic acid, CC: coiled coil, TM: transmembrane. **b**) SDS-PAGE of the co-purified *Pa*FtsWIQBL complex after size-exclusion chromatography. **c**) Western blot showing glycan strand ladders synthesised by divisome core complexes from Lipid II, demonstrating transglycosylase activity. The negative control does not contain any FtsWIQBL (lane 1). WT *P. aeruginosa* and *E. coli* FtsWIQBL complexes (lanes 2 and 4) are active transglycosylases, while the *P. aeruginosa* putative active site mutant FtsW^D275A^IQBL (lane 3) is inactive. **d**) Three representative 2D classes of our *Pa*FtsWIQBL cryo-EM data. **e**) Left panel: side-view of the *Pa*FtsWIQBL cryo-EM density at an overall resolution of 3.7 Å. Protein colours are the same as those in a). Residual density from the detergent micelle is visible around the transmembrane domain in grey. Right panel: model of *Pa*FtsWIQBL, rotated by 120° with respect to the density on the left-hand side. The putative FtsW active site residue D275 is indicated, as is the FtsI active site residue S294. The FtsW loop 219-233 and FtsI loop 45-50 are shown as a dotted line as they were too flexible to build. FtsQ™ and FtsQ^β^ were not resolved and are not shown. **f**) Top view of the periplasmic domain, showing interactions between FtsI, FtsL, FtsB and FtsQ.

### General architecture of the FtsWIQBL complex

All five proteins are resolved in the final cryo-EM reconstruction (Fig. 1d, e), with FtsQ being partially disordered. The density for the membrane domain of *Pa*FtsWIQBL reveals 13 transmembrane (TM) helices, including ten helices from FtsW plus one from each of FtsI, FtsB and FtsL (Fig. S2a). Density for the FtsQ transmembrane helix (FtsQ™) was not observed (Fig. S2b). The detergent micelle density was subtracted from the final reconstruction and the position of the complex in the membrane was approximated using the Orientations of Proteins in Membranes (OMP) webserver^9^ (Fig. 1e, S1b, c).

The periplasmic domains of the *Pa*FtsWIQBL complex extend about 70 Å away from the membrane in a Y-shape, with the FtsI transpeptidase domain (FtsI^TP^) and the FtsQ β-domain (FtsQ^β^) located on opposite arms of the Y, and FtsBL connecting them (Fig. 1e, f). Interestingly, only FtsQ^β^ is well-resolved, while density for the FtsQ polypeptide-transport-associated domain (FtsQ^POTRA^) is only visible in low-resolution maps at high contour level, and density for FtsQ™ is completely absent (Fig. S2d). FtsQ^POTRA^ adopts a slightly different orientation relative to FtsQ^β^ compared to previously determined X-ray structures^10-12^ (Fig. S2d). Taken together, this shows that FtsQ is tethered to FtsWILB only via its FtsQ^β^-FtsB interaction, while FtsQ^POTRA^ and FtsQ™ are flexibly attached in the current complex (Fig. 1f). As FtsQ™ is not visible, we assume it is not in the micelle that contains the other TM segments but might be surrounded by detergent molecules separately. While it has been previously reported that FtsB dimerises and could thus facilitate the dimerisation of core divisome components^13,14^, we find no evidence for higher oligomeric species in our cryo-EM data, and in our current structure dimerisation would be hindered by the presence of FtsI or FtsQ.

FtsL and FtsB have similar folds, each consisting of a long α-helical coiled coil, followed by a short α-helix and a β-strand. Interestingly, the FtsB α-helical coiled coil is interrupted by a small, conserved loop just above FtsB™ that might aid with sterically maintaining the correct insertion depth in the membrane (Fig. 1e, 2a). FtsB and FtsL interact with each other over their entire lengths through mainly hydrophobic interactions, e.g. between FtsL^α1^ and FtsB^α2^ (Fig. 2a and S3a).

**Fig. 2:**
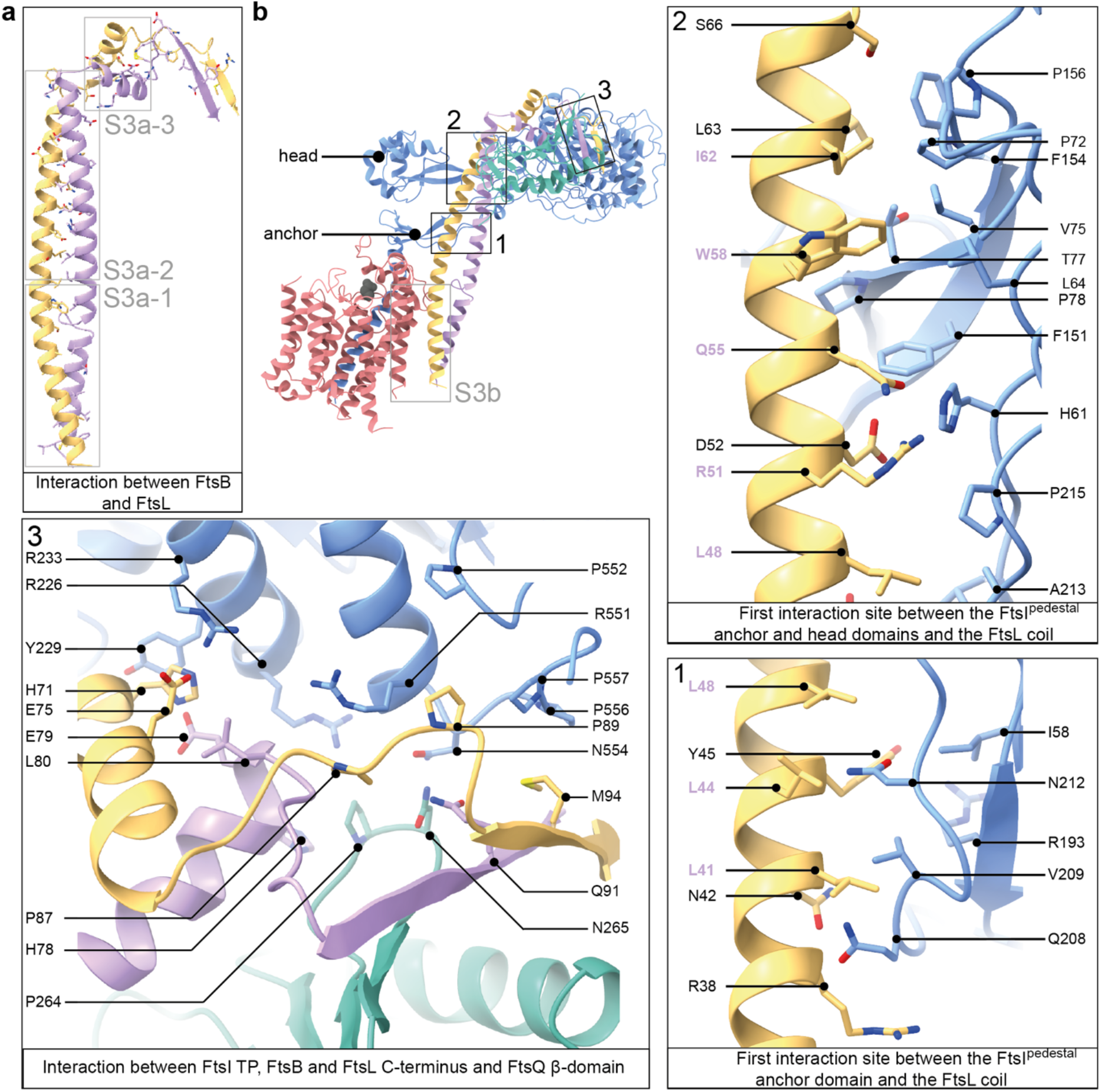
Interactions between FtsI, FtsQ, FtsB, and FtsL. **b**) FtsB and FtsL adopt a similar fold and interact with each other over their whole length. Grey boxes indicate regions that are further discussed in Fig. S3a. **b**) Key interactions within the FtsWIQBL periplasmic domain are highlighted in boxes 1 to 3. The grey box indicates a region that is further discussed in Fig. S3b. Boxes 1-2: the interactions between FtsI^pedestal^ and the FtsL coiled coil are shown in two panels. Most residues in the interface between FtsI and FtsL in this region are hydrophobic or neutral. Residues in FtsL which also face FtsB are highlighted in violet. Box 3: the interaction between FtsI, FtsL, FtsB and FtsQ as seen from the top of the periplasmic domain.

The transglycosylase FtsW and transpeptidase FtsI share two interfaces. The first interface is in the membrane, where FtsI™ interacts with TM8 and TM9 of FtsW – an interaction that closely resembles that of the previously reported RodA-PBP2 elongasome complex from *Thermus thermophilus*^*15*^. The second interaction site is located between the extracellular loop 4 of FtsW (FtsW^EC4^) and the linker between FtsI™ and FtsI^pedestal^. Due to the flexibility of FtsI in this region, not all contacts could be determined unambiguously.

It has been previously reported that the cytoplasmic tail of *E. coli* FtsL is required for the recruitment of FtsW^16^. In the structure presented here, the FtsL cytoplasmic tail could not be traced unambiguously. This could either point towards a transient interaction during recruitment or species-specific differences in the recruitment due to the size of the cytoplasmic tail (11 residues in *P. aeruginosa* vs. 34 residues in *E. coli*). However, we clearly observe FtsL-FtsW interactions in the periplasm (FtsW^EC1^ and FtsL^α1^), and within the membrane through FtsW^TM1^ and the upper three turns of FtsL™. In the latter, the lower part of FtsL™ twists away from FtsW, due to its gyrating coiled-coil interaction with FtsB (Fig. S3b).

### FtsIQBL interactions in the periplasm

FtsL and FtsI form an extensive interface in the periplasm, with a total buried surface area of 1035 Å^2^. The FtsL-FtsI interaction is facilitated by two sites: FtsL^α1^-FtsI^pedestal^, involving anchor and head subdomain residues in FtsI^pedestal^, and FtsL^α2,β1^-FtsI^TP^ (Fig. 2b). Moving along the FtsL coil, the first mainly hydrophobic and neutral interactions occur between FtsL^α1^ (residues L41-L48) and the anchor subdomain of FtsI^pedestal^ (residues I58, R193, Q208-215, R193, Fig. 2b, panel 1). FtsI^pedestal^ slightly wraps around FtsL^α1^, forming a hydrophilic interaction site (FtsI H61 with FtsL R51, D52 and Q55, Fig. 2b panel 2). The final, and mainly hydrophobic, interaction site on FtsL^α1^ involves residues A56-S66 and residues located mainly in the FtsI^pedestal^ head subdomain (L64, P72-P78, F151-P156, Fig. 2b panel 2). Importantly, no direct interaction was observed between the FtsB coiled coil and FtsI (Fig. 2b, S3a).

The second FtsL–FtsI interaction site is located on top of the periplasmic domain (Fig. 2b panel 3): FtsL^α2,β1^-FtsI^TP^. H71 of FtsL^α2^ stacks against Y229 of FtsI^TP^ and is flanked by additional residues in FtsB (E79, L80), FtsI (R233) and FtsL (E75). An additional hydrophobic interface site is formed by several proline residues in both FtsL and FtsI^TP^ [P556, P557 (FtsI) – M94 (FtsL); P557, G478, P477 (FtsI) – P89 (FtsL); R551 (FtsI) -P87 (FtsL)]. Interestingly, FtsB^α3^ and FtsB^β1^ frame a loop in FtsQ between β-strands 11 and 12, forming the only interaction site between FtsI, FtsB and FtsQ [N554 (FtsI) – N265 (FtsQ) – Q91 (FtsB); R226 (FtsI) -P264 (FtsQ); R226 (FtsI) – H78 (FtsB, backbone), R551 (FtsI) – L80 (FtsB, backbone)]. FtsI adopts a structure very similar to previously reported crystal structures^17^, with only minor changes in the FtsI^pedestal^ domain, indicating that FtsL binding does not cause large rearrangements in FtsI^TP^ (Fig. S4a).

Only a few FtsL-FtsQ contacts are present, however FtsL completes an extended β-sheet formed between FtsQ^β^ strands β5 to β12 and FtsB^β1^, by contributing its last β-strand (Fig. 1f, Fig. 2b panel 3). The FtsB-FtsQ interaction recapitulates that of previously determined crystal structures where only small parts of FtsB^11,12^ were resolved (Fig. S4a). In addition, the cryo-EM structure shows an interaction between FtsB^α2^ (starting from E53) and FtsQ^β^ loops (R183, S212-R214, R231). Interruption of the FtsB-Q interface with inhibitors based on the minimal interface could be expected to also disrupt the interface in the context of the divisome core complex^18^.

### Comparison with RodA-PBP2 structures and structure predictions

Cell elongation in rod-shaped bacteria is facilitated by the elongasome that, like the divisome, polymerises and crosslinks PG, but is positioned throughout the cell envelope by MreB filaments^19^. RodA, the elongasome’s transglycosylase is related to FtsW and has previously been structurally characterised using X-ray crystallography both on its own and as a RodA-PBP2 complex^15,20^ (the latter being homologous to FtsWI). The structures of *Pa*FtsW determined here and *Tt*RodA are very similar, with the exception of TM7, which appears to be somewhat flexible in the cryo-EM structure, straighter with respect to that of *Tt*RodA and closer to TM5 than in *Tt*RodA-PBP2 (Fig. S4c). It has been postulated that the movement of TM7 could open a cavity for the binding of the lipid tail of Lipid II to RodA^15^ and the location of TM7 in *Pa*FtsWIQBL creates such a cavity. The putative catalytic residue D275A is located in a deep, highly conserved cleft, as shown in Fig. S4d, that we suggest might harbour the sugar moieties of Lipid II during the transglycosylase reaction.

The most striking observation when comparing *Pa*FtsWIQBL with *Tt*RodA-Pbp2 is the difference in the relative orientations of the TP with respect to the TG domain, despite the fact that the structures of the single proteins superimpose well on their own. Alignment of both complexes on the FtsW/RodA subunits places *Pa*FtsI^TP^ and *Tt*PBP2^TP^ almost opposite to each other, requiring a ∽130° rotation of *Pa*FtsI^TP^/*Tt*PBP2^TP^ for their interconversion (Fig. S5a). The reason for this large difference is unclear, but is possibly caused by the presence of FtsQBL in our divisome structure and the absence of binding partners such as MreCD in the elongasome structure. Alternatively, the differences could be intrinsic to the elongasome and divisome complexes or reflect different, distinct states in the regulatory/catalytic cycle of the enzyme complexes.

We used AlphaFold 2 multimer^21^ (AF2) to predict *Pa*FtsWIQBL and many large-scale and fine features observed in the *Pa*FtsWIQBL structure are predicted correctly by AF2, including the lack of an interaction between the membrane-embedded FtsQ™ and FtsWIBL™. However, in the AF2 model the periplasmic FtsIQBL interaction site is rotated upwards by about 30°, moving FtsI^TP^ closer towards where the peptidoglycan layer is located (Fig. S5a). Furthermore, a small rearrangement in the anchor subdomain FtsI^pedestal^-FtsL^α1^ shifts the interacting residues on FtsI^pedestal^ from 208-212 to 203-206. To understand the implications of these differences, both structures were fitted into a to-scale model of the cell envelope of *E. coli*, produced from a cellular electron cryo-tomogram (Fig. 3a). Using the cryo-EM structure, the active site of FtsI^TP^ does not reach the peptidoglycan layer, but does so in the more extended AF2 model. Since AF2 uses evolutionary couplings between amino acids^21^, in addition to protein structural features that correlate with sequence, it more likely predicts the active state of FtsWIQBL that one would expect to be selected for during evolution, and not the substrate-free form we determined experimentally. Thus, the cryo-EM structure and AF2 prediction may represent the inactive (apo) and active (catalytic) states of the divisome core complex, respectively. Since our sample is active in vitro, we assume that substrate binding by Lipid II might play a key role in the interconversion of the two states. Recently, a study on RodA-PBP2 reported a similar upswinging mechanism^22^, which indicates that the concept of regulating the TP activity via restricting its access to the PG layer might be conserved between the divisome and the elongasome. However, significant differences exist between divisome and elongasome regarding the conformation before the upswinging motion and most likely also between the signals required to initiate this conformational change.

**Fig. 3:**
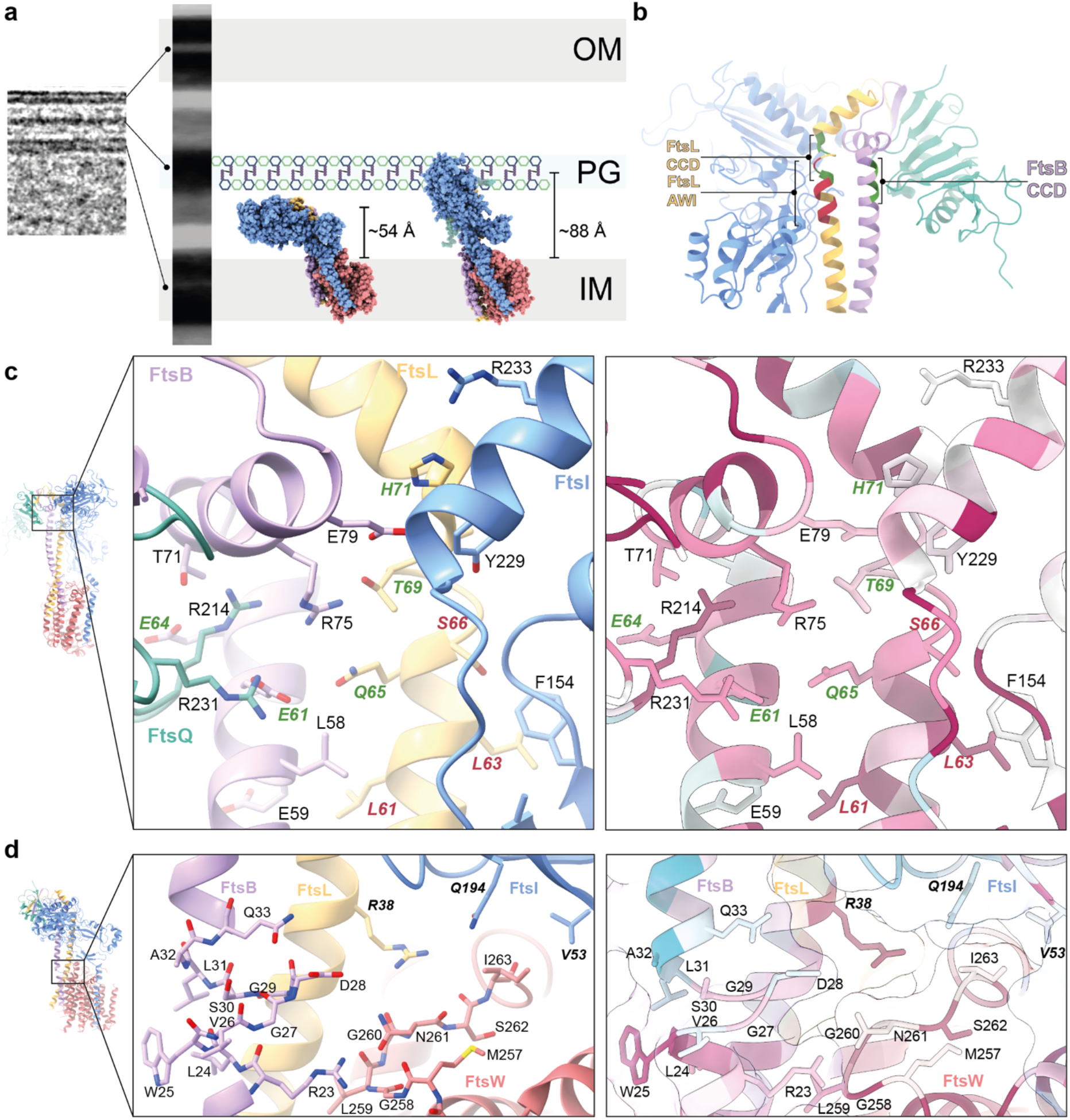
Interactions that affect divisome regulation. **a**) To-scale scheme of the *E. coli* cell envelope generated from a tomogram. The cryo-EM FtsWIQBL atomic model (reported here, left) and the AlphaFold 2 prediction (right) are docked into the inner membrane. The transpeptidase domain of FtsI does not extend to the peptidoglycan layer in our structure, but does so using the AlphaFold 2 structure. The measured distances of the active site residue FtsI^S294^ to the inner membrane plane are indicated. **b**) The C-termini of the FtsB and FtsL coiled coils with highlighted Constriction Control Domain (CCD) and Activation of FtsWI (AWI), showing the FtsB^CCD^ facing FtsQ and FtsL^AWI^ facing FtsI. **c**) Region around the activating mutations FtsL^Q65^ (homologous to E88K in *E. coli*) and FtsB^E61^ (E56 in *E. coli*). Residues with known divisome activating mutations, which lead to smaller cells, are shown in green. Residues with known loss-of-function mutations, which cause a defect in cell division, are shown in red. Sequence conservation analysis (calculated with ConSurf^28^) shows a high degree of conservation for many residues in this area. **d**) Residues surrounding the region of discontinuity in the FtsB coiled coil. This region contains residues with known loss-of-function mutations (shown in bold). Many residues in this region are highly conserved, including FtsL R38 (R61 in *E. coli*), which inserts between FtsI and FtsW and is located close to a highly conserved loop in FtsW (M257-I263, Q279-V285 in *E. coli*) that is in close proximity to the putative active site residue D275 (D297 in *E. coli*).

FtsN has been reported to trigger constriction in cells^23,24^. It is the last protein to be recruited to the division site in *E. coli* and its recruitment is dependent on the presence of earlier divisome proteins, including FtsA, FtsQ and FtsI^23,25-27^. However, we have not been able to generate a biochemically-stable *E. coli* FtsN-FtsWIQLB complex and previous studies reported that the addition of the FtsN periplasmic domain did not yield an increase in *P. aeruginosa* TG activity *in vitro*^*7*^. Whether FtsN activates the core divisome beyond the TG activity levels seen here through binding of FtsQLB, or whether the *in vitro* sample cannot be further activated will need further investigations including addition of other divisome components, e.g. FtsA, FtsN and/or DedD, as well as the substrate Lipid II. It is also possible that FtsN is involved in regulating TP activity.

### Interactions that affect divisome regulation

The Constriction Control Domain (CCD) of the divisome was identified previously from a set of mutations that allow partial or complete bypass of the requirement for FtsN in *E. coli*^29,30^. In our structure these residues cluster at the top of FtsL^α1^ and FtsB^α2^, with the FtsB CCD mutations facing FtsQ^β^. Furthermore, in close proximity are the Activation of FtsWI (AWI) residues on FtsL^α1^ that display a dominant-negative phenotype when mutated^16^ (Fig. 3b).

The CCD residues of FtsB^E56^ (*Ec*E61G/A/K/V/H) and FtsL^Q65^ (*Ec*E88K/V) point inwards into a positively charged cavity formed by three arginine residues (FtsQ^R214^, FtsQ^R231^, FtsB^R75^), flanked by FtsL^T69^ (*Ec*D93G, CCD residue) on one side and FtsB^E64^ (*Ec*D59V, CCD residue) and FtsB^T71^ on the other side. This interface contains many charged and highly conserved residues (Fig. 3c), and removal of a charge or introduction of the opposite charge could well result in destabilisation of the interface and potentially increased flexibility of the protein. This may allow FtsWIQBL to more readily adopt an elongated, a more active conformation, as possibly indicated by the AF2 model, and with less or no activation signal, for example from FtsN.

A dominant negative phenotype was previously reported for the AWI mutation *Ec*L86F^16^ and its *P. aeruginosa* equivalent FtsL^L63^ interacts with FtsI^F154^ in FtsI^pedestal^ (Fig. 3c). Replacing the leucine with the bulkier phenylalanine likely causes a steric clash that weakens the FtsI-FtsL interaction. FtsL^S66^ (*Ec*N89S, *Pa*S66D) was classified as a CCD mutation in *E. coli*^*29*^, but has a dominant negative phenotype in *P. aeruginosa*, with reduced *in vitro* TG activity^7^, likely due to a steric clash with FtsI^F154^. Mutation of FtsL^L61^ (*Ec*84K/D) has a dominant-negative phenotype and affects FtsL localisation to the septum^16,31^, indicating that the integrity of the FtsBL coiled-coil interaction is vital for a functional complex, as is well supported by our structure where the FtsBL coiled coil is at the centre of the complex.

The AWI residue FtsL^R38^ (*Ec*R61C) is highly conserved and located between FtsW^M257-I263^ and the anchor subdomain of FtsI^pedestal^ (Fig. 3d). Mutation of this residue causes a dominant-negative phenotype in both *E. coli* and *P. aeruginosa* and reduced *in vitro* TG activity in *P. aeruginosa*^*7,16*^. FtsL^R38^ might interact with the highly conserved FtsW^G260^ and FtsW^S262^ residues and stabilise FtsW^M257-I263^ together with FtsB^R23^, a hypothesis supported by the fact that the corresponding loop in RodA and RodA-PBP2 is disordered^15,20^. The aforementioned FtsB coiled-coil discontinuity, C-terminal of FtsB^R23^, might be required to allow for some flexibility of FtsB during the catalytic cycle of FtsW. *Ec*FtsI^L62P^ and *Tt*PBP2^L43R^ correspond to *Pa*FtsI^V53^ and result in a strong cell division defect and reduced TG activity *in vitro*, respectively^15,32^. *Pa*FtsI^V53^ interacts with *Pa*FtsW^I257^ in our structure, bringing the linker between FtsI™ and FtsI^pedestal^ in close proximity to FtsW. Thus, FtsW^M257-I263^ presents an interaction site for FtsL, FtsB and FtsI in close proximity to the putative FtsW active site.

The recruitment of FtsQ to midcell requires FtsK^10,26,33^. AF2 predicts that the interaction between FtsK^1-222^ and FtsWIQBL, occurs through FtsQ^POTRA^ β2 and α3; this interaction is also identified by coevolutionary coupling analysis using EVcouplings^34^ as an FtsQ-FtsK interaction hotspot (6/10 couplings, Fig. S6a, b). Residues previously identified as impairing FtsK recruitment when mutated^10^ (*Ec*Q108, *Ec*V92, *Ec*V111, *Ec*K113) map onto this conserved region (Fig. S6c). We copurified an FtsQK^1-222^ complex using *E. coli* proteins, which confirms that the two proteins interact tightly and constitutively (Fig. S6d). To further our understanding of divisome recruitment and regulation, in the future larger divisome complexes will need to be assembled. For example, divisome interactions with FtsA, through FtsN, FtsK and possibly FtsQ have the potential to modify the conformations, oligomeric states and activities of the core complex and its enzymes.

We report the structure of the essential bacterial cell division complex and important antibiotic target FtsWIQBL from *Pseudomonas aeruginosa* and show that *Pa*FtsWIQBL forms a stable Y-shaped complex that harbours intrinsic TG activity. Our *Pa*FtsWIQBL structure is able to explain many subunit contacts that have previously been shown to be important through loss-of-function and bypass mutations. In addition, the generated AF2 model reveals a different, likely catalytically competent state that allows for peptidoglycan crosslinking by FtsI. While our analysis hints at the nature of the catalytic state, further research is needed to resolve more states and their associated conformation changes, which possibly requires the addition of activating proteins such as FtsA and/or FtsN as well as the substrate Lipid II and its derivatives or products as ligands. It will be particularly exciting to resolve the enzymatic mechanism of the FtsW TG since it is a very promising drug target for novel antibiotics. To gain a deeper understanding of the mechanism of the divisome, the inclusion of upstream and downstream proteins e.g., FtsEX, FtsK, PBP1b, and DamX will also be necessary. Our work is an important milestone in the 25-year quest for a molecular understanding of the ancient, near ubiquitous and medically important process of FtsZ-based bacterial cell division.

## Acknowledgements

We thank all members of the MRC LMB electron microscopy facility for excellent EM support and T. Darling and J. Grimmett for computing support. We are grateful to Dougal Ritson and Meng Su for their help and advice during the Lipid II preparation and to Holger Kramer (1979-2022) for LC-MS and MALDI-TOF of Lipid II samples (all MRC LMB). N-terminal His-tagged *Sa*Pbp4(aa21-383) was a kind gift of the Walker lab (Harvard, MA). We thank Tanmay Bharat and Stephan Tetter for their feedback on the manuscript. This work was funded by the Medical Research Council (U105184326) and the Volkswagen Stiftung “Life?” programme (both to J.L.).

## Author Contributions

L.K., F.v.d.E and M. J. purified the *Pa*FtsQBLWI complex. L.K. collected and processed the cryo-EM data. F.v.d.E and M. J. purified the *Ec*FtsQBLWI complex. F.v.d.E performed the FtsWIQBL activity assays. N.J. cloned and purified the FtsQ-FtsK complex. V.H. prepared, measured and processed the tomogram. J.L. provided the concept and L.K., M. J. and J.L. prepared the manuscript. All authors contributed to the materials and methods section and to editing the manuscript.

## Additional Information

Supplementary Information is available for this paper.

Correspondence and requests for materials should be addressed to Jan Löwe.

## Data Availability

The final cryo EM map has been deposited in the Electron Microscopy Data Bank (EMDB) with the accession code EMD-16042. The final model has been deposited with the Protein Data Bank (PDB) with the accession code 8BH1.

## Materials and methods

### Cloning

#### P. aeruginosa FtsWIQLB

Cloning, expression and purification of *Pa*FtsWIQBL (Uniprot: Q9HW00 (FtsW), G3XD46 (FtsI), G3XDA7 (FtsQ), Q9HVZ6 (FtsL), Q9HXZ6 (FtsB)) were adapted from a previously published protocol^1^. FtsW and FtsI were expressed on a pET-Cola vector, and FtsQ, FtsL, and FtsB were expressed on a pET-Duet vector. The genes encoding *P. aeruginosa* FtsI and FtsL were obtained by PCR from the strain PAO1 using primers PaFtsI.fwd/rev and PaFtsL.fwd/rev respectively (primers: Table S1). The codon-optimised sequences for FtsB-His^6^, His_6_-SUMO-FtsQ, and His_6_-SUMO-FLAG-FtsW were ordered as gBlocks (IDT). These fragments were extended by PCR to generate overhangs for Gibson assembly using primers PaFtsQB.fwd/rev and SUMO-PaFtsW.fwd/rev, respectively. The FtsI, FtsW and pET-Cola (amplified using primers pET-Cola.fwd/rev) fragments were assembled by Gibson assembly forming the plasmid pLK2. Later, the FLAG-tag was exchanged for a Twin-Strep-tag using PCR with primers Strep-PaFtsWI.fwd/rev and blunt-end ligation (plasmid pLK4). The mutant FtsW^D275A^ was generated from the pLK4 plasmid by site directed mutagenesis using primers PaFtsW_D275A.fwd/rev. The pET-Duet plasmid carrying FtsQ, FtsL, and FtsB was generated in two consecutive Gibson assembly reactions. First, the extended FtsQ, FtsB and pET-Duet (amplified using pET-Duet_FtsQB.fwd/rev) fragments were assembled by Gibson assembly and the generated plasmid was isolated, and opened with primer pET-Duet_FtsL.fwd/rev. Second, the opened plasmid and extended FtsL fragments were assembled by Gibson assembly forming plasmid pLK1. The DNA for the His-tagged catalytic subunit of *S. cerevisiae* SUMO protease Ulp1 (residues 403-612) was ordered as a gBlock (IDT) and introduced into a pBad vector using Gibson assembly forming the plasmid pLK3. pLK3 contains an arabinose inducible promoter, a chloramphenicol resistance gene and a p15A origin. All constructs were confirmed by sequencing.

#### E. coli FtsWIQLB

The *E. coli* FtsWIQLB complex (Uniprot: P0ABG4 (FtsW), P0AD68 (FtsI), P06136 (FtsQ), P0AEN4 (FtsL), P0A6S5 (FtsB)) was expressed in insect cells through a single baculovirus vector, which was assembled using the biGBac system^2^. To ensure equal expression levels, a fusion of FtsW and FtsI, as is found naturally in some organisms (data not shown), was created with a GSGASG cytoplasmic linker between the FtsW C-terminus and FtsI N-terminus. All genes were ordered as gBlocks (IDT). An NdeI site was introduced downstream of the BamHI site of pLIB (AddGene) by site-directed-mutagenesis (SDM) resulting in pFE661. The *E. coli* FtsI gene, with overhangs for Gibson assembly, was introduced into pLIB using Gibson assembly forming pFE668. The *E. coli* FtsW gene, with an N-terminal Twin-Strep-tag and NdeI site, was amplified using primers MTAstrep.for and WlinkerI.rev and introduced into pFE668 by Gibson assembly. The gene expression cassette (GEC), containing the polyhedron promoter and Twin-Strep-FtsW-GSGASG-FtsI coding sequence, was amplified using primers casI.for and casv.rev^2^ and introduced into SwaI-opened pBig1b (Addgene) by Gibson assembly, resulting in pFE756. The *E. coli* FtsQ gene was cloned into NdeI/HindIII of pLIB, resulting in pFE686. The *E. coli* FtsB gene was amplified using primers EcB_acebac1.for/rev and introduced into pACEBac1 (Geneva Biotech) by Gibson assembly, resulting in pFE658. The *E. coli* FtsL gene, with an N-terminal 10xHis site followed by a TEV site, was introduced into pLIB by Gibson assembly forming pFE674. To increase expression levels of FtsL, the mutations S3N, R4K and V5L were introduced by SDM using primers Ltripple.for/rev^3^. The GECs of FtsQ, FtsB and His_10_-TEV-FtsL were amplified by PCR using casI.for/rev, casII.for/rev and casIII.for/rev, respectively and introduced into SwaI-opened pBig1a by Gibson assembly, resulting in pFE749. In a final Gibson assembly reaction the PmeI fragments of pFE756 (Twin-Strep-FtsW-FtsI in pBig1b) and pFE749 (FtsQ-FtsB-10xHisFtsL in pBig1a) were combined with PmeI-opened pBIG2ab resulting in pFE758. All constructs were confirmed by sequencing.

#### E. coli FtsQK^1-122^

The genes encoding *E. coli* FtsQ-His_6_ (Uniprot P0613) and TwinStrep-FtsK^1-222^ (Uniprot P46889) were ordered as gBlocks (IDT) and cloned into pLIB using Gibson assembly. The GECs were amplified and inserted into pBIG1a^2^ forming pNJ069.

### Baculovirus generation

Baculoviruses were created containing pFE758 and pNJ069 for insect cell expression of *E. coli* FtsWIQLB and FtsQK^1-222^ respectively. Recombinant baculoviral genomes were generated by TN7 transposition in DH10bacY cells^4^. This bacmid was used to transfect Sf9 cells (Thermo Scientific) using FuGENE (Promega). After 3-5 days, the culture was centrifuged and the virus-containing supernatant harvested and stored till use with 1% fetal bovine serum (FBS) added.

### Bacterial expression

*P. aeruginosa* FtsWIQLB was expressed in *E. coli* cells. pLK1, pLK2 and pLK3 were sequentially transformed into *E. coli* C43(DE3) cells. 120 mL of overnight culture were added to 12 L TB media with 2 mM MgCl_2_, containing kanamycin (25 μg/mL), chloramphenicol (25 μg/mL) and ampicillin (50 μg/mL), and grown at 37°C to an OD_600_ of 0.7. Protein expression was induced with 1 mM IPTG and 1 g arabinose/L and continued at 18°C overnight. Cells were harvested by centrifugation for 20 min at 4,000 rpm and 4°C, then flash frozen in liquid nitrogen and stored at −80°C.

### Insect cell expression

*E. coli* FtsWIQLB and FtsQK^1-222^ were expressed in insect cells. Sf9 cells were grown in Insect-Xpress medium (Lonza) to a density of 1.5-2 million cells/ml. They were infected with ∽1% amplified baculovirus and harvested by centrifugation after 60-70 hrs, at a cell viability of ∽80%. Cell pellets were washed with phosphate buffered saline (PBS), flash frozen in liquid nitrogen and stored at −80°C.

### Protein purifications

All purifications were done at 4°C.

#### P. aeruginosa FtsWIQBL

Cells from 12 L of *E. coli* culture were resuspended in a final volume of 300 ml Lysis Buffer (20 mM HEPES, 150 mM NaCl, 20 mM MgCl_2_, pH 7.5, 1 mM DTT) containing DNase (Sigma) and RNase (Sigma) and passed through a cell disruptor (Constant Systems) at 25,000 psi twice. Subsequently, the lysate was centrifuged for 1 hr at 4°C and 45,000 rpm (Type 45 Ti rotor, Beckmann). The membranes were homogenised using a dounce tissue grinder (Whaeton) in Solubilisation Buffer (20 mM HEPES, 500 mM NaCl, 20 % glycerol, pH 7.0). Lauryl maltose neopentyl glycol detergent (LMNG, Anatrace) was added to a final concentration of 1% (w/v) to solubilise the membrane-bound proteins while rotating for at least 1 hour at 4°C. The volume was doubled with Solubilisation Buffer and 12 μl benzonase (Merck) were added. The solution was centrifuged for 1 h at 4°C and 45,000 rpm (Type 45 Ti rotor, Beckman) and the supernatant was bound to 2 mL (CV) of equilibrated Strep XT4 Flow beads (IBA) while rotating at 4°C for 2 h. Beads were collected in a column and washed with 50 mL of Wash Buffer 1 (20 mM HEPES, 500 mM NaCl, 20 % glycerol, pH 7.0, 0.1% LMNG) and 50 ml of Wash Buffer 2 (20 mM HEPES, 500 mM NaCl, 20% glycerol, pH 7.0, 0.01% LMNG). Protein was eluted from the Strep beads over 10 CV in 2 mL fractions with Elution Buffer (20 mM HEPES, 500 mM NaCl, 20% glycerol, 50 mM biotin, pH 7.0, 0.005% LMNG). All elution fractions were pooled and concentrated to 50 μL using 100 kDa centrifugal concentrators (Vivaspin). The concentrated sample was further purified using a Superose 6 Increase 3.2/300 size-exclusion column (Cytiva), equilibrated in Size Exclusion Buffer (20 mM HEPES, 300 mM NaCl, pH 7.0, 0.005% LMNG). Fractions were either used directly for cryoEM grid preparation or flash frozen in liquid nitrogen, before being stored at −80°C. For the activity assay (see below) protein concentration was determined using Bio-Rad Protein Assay dye reagent concentrate (Bio-Rad).

#### E. coli FtsWIQBL

Sf9 Cells were resuspended in Lysis Buffer (20 mM HEPES pH 8.0, 500 mM NaCl, 10% glycerol, 2 mM TCEP), containing 1 mM PMSF, protease inhibitor tablets (cOmplete EDTA-free PI (Roche), 1 per 25 mL), DNase (Sigma), RNase (Sigma) and sonicated for 2 min (1 s pulse on, 10 s pulse off, 70% intensity), after which 2 mM EDTA was added. The lysate was centrifuged for 1 h at 235,000 xg. The pellet was homogenised using a dounce tissue grinder (Whaeton) in Solubilisation Buffer (20 mM HEPES pH 8.0, 10 mM MgCl_2_, 500 mM NaCl, 20% glycerol), containing 1 mM PMSF, and PI tablets. Detergent glyco-diosgenin (GDN, Anatrace) was added to a final concentration of 1% to solubilise the membrane-bound proteins while rotating at 4°C for 2 h. The mixture was centrifuged for 10 min at 3,200 xg, 4°C to remove nuclei. The supernatant was 5-fold diluted with Strep Buffer (20 mM HEPES, pH 8.0, 350 mM NaCl, 10% glycerol, 1 mM TCEP) in the presence of Benzonase (Merck) and the NaCl concentration was brought down to 350 mM before centrifuging for 45 min at 142,000 g. The supernatant was recycled over a 5 mL StrepTrap-HP column (Cytiva) overnight. The StrepTrap-HP column was washed with 50 column volumes of Strep Buffer including 0.01 % GDN at 5 mL/min. The protein complex was eluted in Strep Buffer supplemented with 2.5 mM desthiobiotin (Sigma) at 1 mL/min. Fractions containing the complex were pooled and concentrated to 50 μL using 100 kDA centrifugal concentrators (Vivaspin) and further purified using a Superose 6 Increase 3.2/30 size-exclusion column (Cytiva) in SEC buffer (20 mM HEPES pH 8.0, 350 mM NaCl, 10% glycerol, 1 mM TCEP, 0.01% GDN). Protein concentration was determined using Bio-Rad Protein Assay dye reagent concentrate (Bio-Rad) and fractions were aliquoted, then flash frozen in liquid nitrogen before being stored at −80°C.

#### E. coli FtsQK

Cells from 1 L of culture were resuspended in 50 mL Lysis Buffer (50 mM Tris, 500 mM NaCl, pH 8.0) supplemented with DNase (Sigma), RNase (Sigma), and protease inhibitor tablets (cOmplete EDTA-Free PI (Roche), 1 per 25 mL) and sonicated for 2 min (1 sec on, 10 sec off, 70% intensity). The lysate was centrifuged for 25 min at 4°C and 25,000 g (25.50 rotor, Beckmann) and its supernatant subsequently for 1 h at 4°C and 200,000 g (Ti 45 rotor, Beckmann). The membranes were homogenised with a Dounce homogeniser and solubilised in 35 mL Solubilisation Buffer (50 mM Tris, 350 mM NaCl, pH 8.0, 1% GDN, 10% glycerol) for 2 h at 4°C. The solution was diluted to 50 mL with Dilution Buffer (50 mM Tris, 350 mM NaCl, pH 8.0, 10% glycerol) and centrifuged for 30 min at 4°C and 80,000 g (Ti 75 rotor, Beckmann). The supernatant was diluted to 500 mL with dilution buffer and recycled overnight over a 1 mL StrepTrap-HP column (Cytiva). The column was washed with 70 mL of buffer A1 (50 mM Tris, 350 mM NaCl, 0.006% GDN, 10% glycerol, pH 8.0) and the complex was eluted with 20 mL of Buffer A2 (50 mM Tris, 350 mM NaCl, 0.006% GDN, 10% glycerol, 2.5 mM desthiobiotin, pH 8.0) in 2 mL fractions, and fractions containing FtsQK were pooled. This eluate was bound to an equilibrated 1 mL HisTrap (Cytiva) column and eluted using a step gradient from Buffer B1 (50 mM Tris, 350 mM NaCl, 0.006% GDN, 10% Glycerol, 20 mM imidazole, pH 8.0) to Buffer B2 (50 mM Tris, 350 mM NaCl, 0.006% GDN, 10% glycerol, 1 M imidazole, pH 8). The HisTrap elution was concentrated to 50 μL using a 100 kDa cutoff centrifugal concentrator (Vivaspin). The complex was purified further using a Superose 6 Increase 3.2/300 size-exclusion column in SEC buffer (50 mM Tris, 100 mM NaCl, 0.006% GDN, 10% glycerol, pH 8.0).

#### S. aureus PBP4

His-tagged PBP4^21-383^ from *Staphylococcus aureus* (*Sa*Pbp4) was expressed as described previously^5^ and purified as follows. Cells were lysed in Buffer A (50 mM Tris pH 7.5, 500 mM NaCl) containing 1 mM PMSF, DNase, RNase and PI tablets using a cell disruptor (Constant Systems) at 25 kpsi. The lysate was centrifuged at 100,000 g for 30 min. The supernatant was supplemented with 1% Buffer B (Buffer A + 1 M imidazole, pH 7.5) and recycled twice over a 5 mL HisTrap-HP column (Cytiva). The column was washed with 150 mL 1% B, 550 mL 2% B and eluted in steps of 5%, 30% and 50% B. Fractions containing *Sa*Pbp4 were concentrated by centrifugal filtration (Vivaspin) and further purified over a Superdex 200 PG 16/60 size-exclusion column (Cytiva) in SEC buffer (20 mM MES pH 6.0, 300 mM NaCl). The fractions of the monomer peak were combined and concentrated to 52 g/L by centrifugal filtration (Vivaspin), then flash frozen in aliquots before being stored at −80°C.

### Cryo-EM single particle structure determination

#### Grid preparation

Grids were prepared with freshly purified protein (*Pa*FtsWIQBL) from the peak fraction of the SEC elution at a final concentration of 1.2-1.3 g/L, diluting with SEC buffer if necessary. 3 μL of sample were pipetted onto a freshly glow discharged (PELCO easiGlow, 25 mA, 45 sec) 300 mesh Cu 0.6/1 grid (Quantifoil) and blotted for 4 s at strength 4, 100% humidity and 4°C, before plunge-freezing in liquid ethane using a Vitrobot Mark IV (Thermo Fisher Scientific, TFS).

#### Data collection

Data from 5531 micrographs was collected on a Titan Krios G3 (TFS) equipped with a Gatan K3 camera and a Gatan Quantum energy filter (20 eV slit width). TFS’s EPU software was used to collect the micrographs with fringe-free imaging in counting mode at a nominal pixel size of 1.09 Å. The exposure time was 2.5 sec at a dose of 21 e/px/s, defocus between −1.2 and −3 μm, and 40 fractions per micrograph were collected.

#### Data processing

Unless stated otherwise, all processing was done in RELION 4.0^6^. Motion correction was performed using RELION’s own implementation of the MotionCor2 algorithm^7^ with 5×5 patches. Subsequently, CTFFind-4.1^8^ was used for CTF estimation. The initial reference was generated in CryoSPARC^9^ from a different dataset of the same sample (not used for the final reconstruction). Particles were picked with Topaz^10^ 7,276,623 particles total, (1,398 per micrograph on average) and extracted 4x binned with a boxsize of 70 px and a pixel size of 4.36 Å/px. The particles were split into seven subsets for initial 3D classification into three classes. The best classes of each job were selected and refined. The refined particles were combined into two sets of particles and subjected to 3D classification without alignment. The best classes from each alignment were selected (1,997,326 and 2,922,553 particles total), combined and subjected to a 3D refinement. Subsequently, a mask (extended by 2 px and added soft edge of 2 px) was used to subtract the micelle density from the complex. The subtracted particles were subjected to a 3D classification without alignment and two classes were selected (344,300 and 461,499 particles), re-extracted (2x binned with a boxsize of 140 pix and a pixel size of 2.18 Å/px) and subjected to 3D refinement. A 3D classification without alignment was run and the best class (160,615 particles) was selected. The particles were re-extracted (no binning, 280 pix, 1.09 Å/px). The particles were subjected to one round of polishing, 3D refinement, CTF refinement and 3D refinement. The micelle was subtracted from the particles and the particles were subjected to a round of 3D refinement and 3D classification without alignments. From this 3D classification, the best class was selected (136,364 particles) and 3D refined two times, the second time using a mask that excluded the POTRA domain of FtsQ. After postprocessing with the calibrated pixel size (see below) the final reconstruction had an overall resolution of 3.7 Å as determined by Fourier shell correlation (FSC, cutoff 0.143).

#### Model building

The pixel size was calibrated using Chimera^11^ and a published crystal structure of *Pa*FtsI (PDB:3OCN). The pixel size was adjusted to 1.05 Å/px during postprocessing in RELION. This map was used to fit a model of *Pa*FtsWIQBL predicted with AlphaFold2^12,13^ and manually adjusted using MAIN^14^ and Coot^15^ (Version: 0.9.8.3), and real-space refined using Phenix^16^ (Version: 1.19.2-4158). Figures of the structure were prepared using ChimeraX-1.4^17^.

### Lipid II extraction

Lipid II was extracted from *Enterococcus faecalis* (DSMZ 2570) as described before^18^. Briefly, an overnight culture of *E. faecalis* was diluted 1:100 into 3 L of Brain heart infusion (BHI, Merck) and grown at 37°C, 180 rpm to an OD_600_ of 0.7. Vancomycin (Sigma) and moenomycin (Santa Cruz) were added at 10 μg/mL and 5 μg/mL, respectively, and cells were centrifuged 20 min later in pre-cooled bottles at 4,500 g for 20 min. The cell pellets were resuspended in BHI, spun again in Falcon tubes at 3,200 g for 10 min, flash frozen in liquid nitrogen and stored at −20°C overnight. Frozen cell pellets were thawed in a total of 30 mL phosphate-buffered saline (PBS), divided equally into two glass 250 mL Erlenmeyer flasks and 17.5 mL chloroform and 35 mL methanol were added. After 2 h of vigorously stirring at room temperature, the mixture was spun in Teflon tubes for 10 min at 4,000 g at 4°C and the supernatant from each Erlenmeyer was combined with 30 mL chloroform and 22.5 mL PBS. The mixture was stirred vigorously for 2 h at room temperature, then spun for 10 min at 4,000 g at 4°C. The tubes were left at room temperature for 1 h until the supernatant was clear. The interface was then transferred with a glass Pasteur pipette to a 25 mL separatory funnel and left to settle overnight at 4°C. The lower organic phase was discarded and the interface dried in a 25 mL round bottom flask on a rotary evaporator at 40°C. The dried interface was resuspended in 7.5 mL pyridinium acetate:n-butanol (1:2) (PB) and 7.5 mL n-butanol-saturated water. Pyridinium acetate was previously prepared by adding 51.5 mL glacial acetic acid dropwise to 48.5 mL pyridine and filtered before use. The Lipid II extract was transferred to a 25 mL separatory funnel and the bottom phase was re-extracted with 5 mL PB. The top phase from the re-extraction was added to the top phase in the separatory funnel and extracted three times with 5 mL n-butanol-saturated water. The top phase was dried using a rotary evaporator at 40°C and resuspended in chloroform:methanol (1:1), partially dried under a stream of nitrogen gas and then transferred to a 250 μL non-stick glass vial (Agilent), in which it was dried completely. This was repeated 4 times to ensure efficient transfer before resuspending the Lipid II extract in 210 μL chloroform:methanol (1:1). The Lipid II extract was assessed by spotting 1-2 μL on a HPTLC silica gel 60F254 plate (Merck). The TLC plate was developed in a mixture of chloroform:methanol:water:ammonia (88:48:10:1) and Lipid II was visualised by heating the plate after soaking in phosphomolybdic acid (PMA) as described previously^19^.

### Deprotection of FMOC-BDL

D-Lys-Biotin (BDL) was prepared following a standard deprotection protocol^5^. Briefly, 15 mg Fmoc-D-Lys(Biotin)-OH (Santa Cruz Biotechnology) was stirred in 3.1 mL of 20% piperidine/dimethylformamide and 466 μL toluene for 40 min at room temperature, then dried in vacuum at 50°C. The sample was resuspended in 5 mL water, stirred for 2 h at room temperature and then filtered through a 0.22 μm filter. The filtrate was pipetted into a tared tube, frozen on dry ice and then freeze dried. The residue was dissolved in water to make a 10 mM stock, aliquoted and stored at −20°C.

### Transglycosylase activity assay

Lipid II used to monitor glycosyltransferase activity of the protein complex was dried using a nitrogen stream and dissolved in an equal volume of dimethyl sulfoxide (DMSO). The reaction and detection of glycan strands was adopted from previously published protocols^5,20-22^. Briefly, *Pa*FtsWIQLB and *Ec*FtsWIQLB were mixed at a final protein concentration of 1 μM in 10μL with 1 μL 10x Reaction Buffer (500 mM Tris pH 7.5, 200 mM MnCl_2_), 1 μL DMSO and 1 μL Lipid II and incubated for 30 min at 25°C. Proteins were heat inactivated for 2 min at 95°C. Lipid II and glycan strands were labelled by incubating the reaction with 26 μM *Sa*Pbp4 and 20 mM BDL for 1 h at 25°C. An equal volume of Laemmli SDS-PAGE buffer was added and the mixtures were heat inactivated for 3 min at 95°C. Glycan strands were separated from Lipid II on a 4-20% Criterion TGX polyacrylamide gel (Bio Rad), run for 45 min at 200 V. After blotting onto PVDF membrane, the blot was incubated for 2 h in Superblock blocking buffer TBS (Thermo Scientific), followed by incubation with a 1:5000 dilution of IRDye800CW Streptavidin (LI_COR Bioscience) in TBS buffer at room temperature for 1 hr. The blot was washed three times in PBS buffer and bands were visualised using an Odyssey CLx imaging system (Li-COR Bioscience).

### Electron cryo-tomography of *E. coli* cells

A culture of *E. coli* strain B/r H266 expressing plasmid pRBJ212^23^ were grown in LB media at 37°C, 180 rpm to an OD_600_ of 0.6. Cells were concentrated 10x by centrifugation and mixed with 10 nm protein A gold fiducials in a 1:10 ratio. 4 μL of this mixture was applied to 200-mesh copper grids with a Quantifoil R2/2 support film, back-blotted and plunge-frozen in liquid ethane using a manual plunger. Cells were thinned for electron cryo-tomography by cryo-focused ion beam milling (cryoFIB) using a Scios dual beam FIB-SEM (FEI). Before milling, grids were sputtered with a protective layer of organic platinum using the gas injection system. Lamellae were milled in a stepwise fashion, gradually reducing the beam current as the lamellae were thinned, starting at 1 nA and polishing at 30 pA and at a nominal milling angle of 10°. CryoET was carried out on a Krios microscope (ThermoFisher) equipped with a Gatan imaging filter and K2 camera. Tilt series were collected using serialEM software^24^, using a bidirectional tilt scheme from −10° (to flatten the lamella) with a 2° increment and a total dose of 112 e^-^/Å^2^, divided over 56 images, each with 10 frames. The pixel size was 3.7 Å and the defocus target was −5 μm. Frame alignment and tilt series alignment were performed using IMOD^25^ and a 2x binned aligned tilt series was generated. The aligned tilt series were used to generate a SIRT reconstruction using tomo3D^26^, which was then low-pass filtered to 20 Å.

## Supplementary figures

**Table S1:**
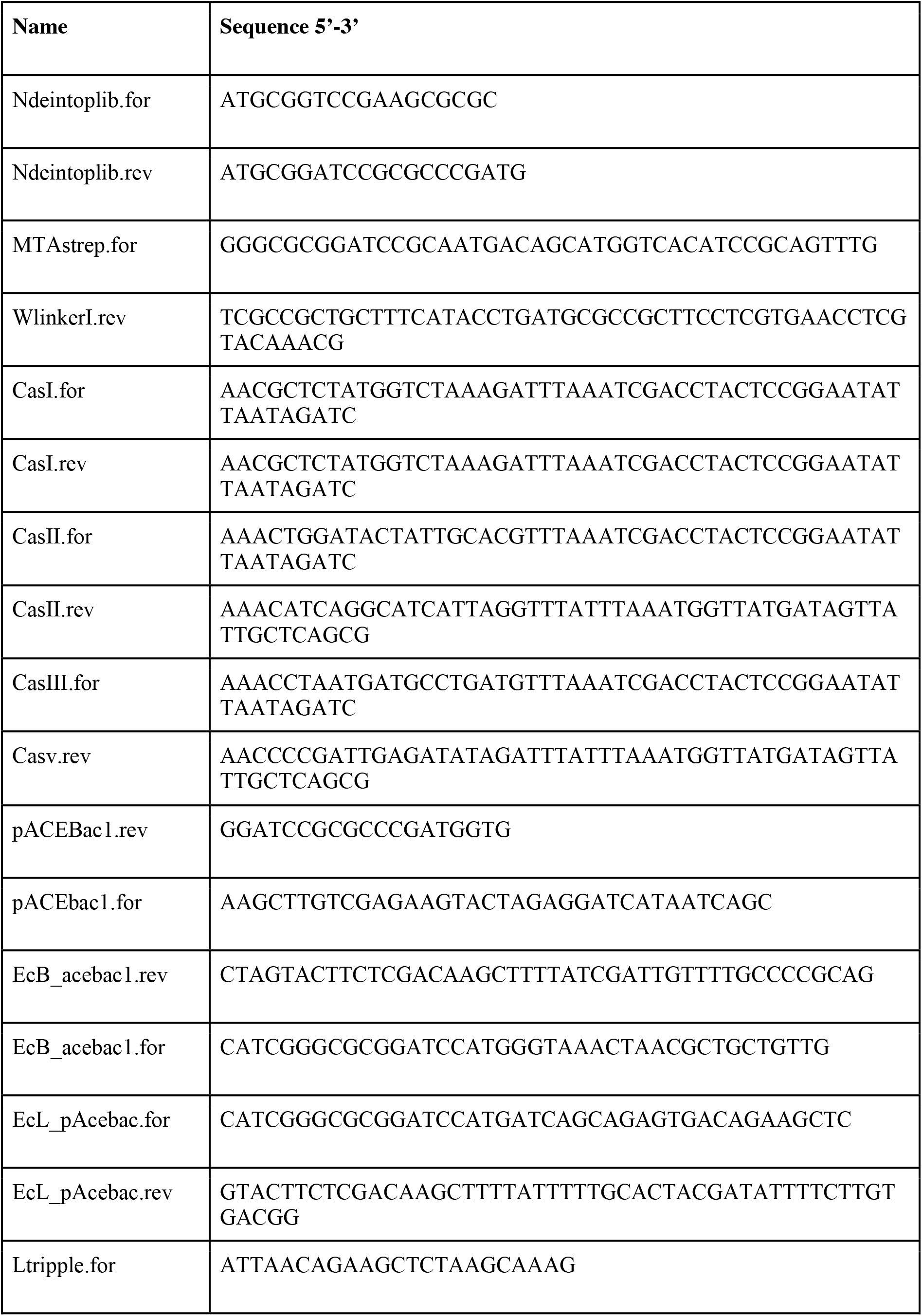
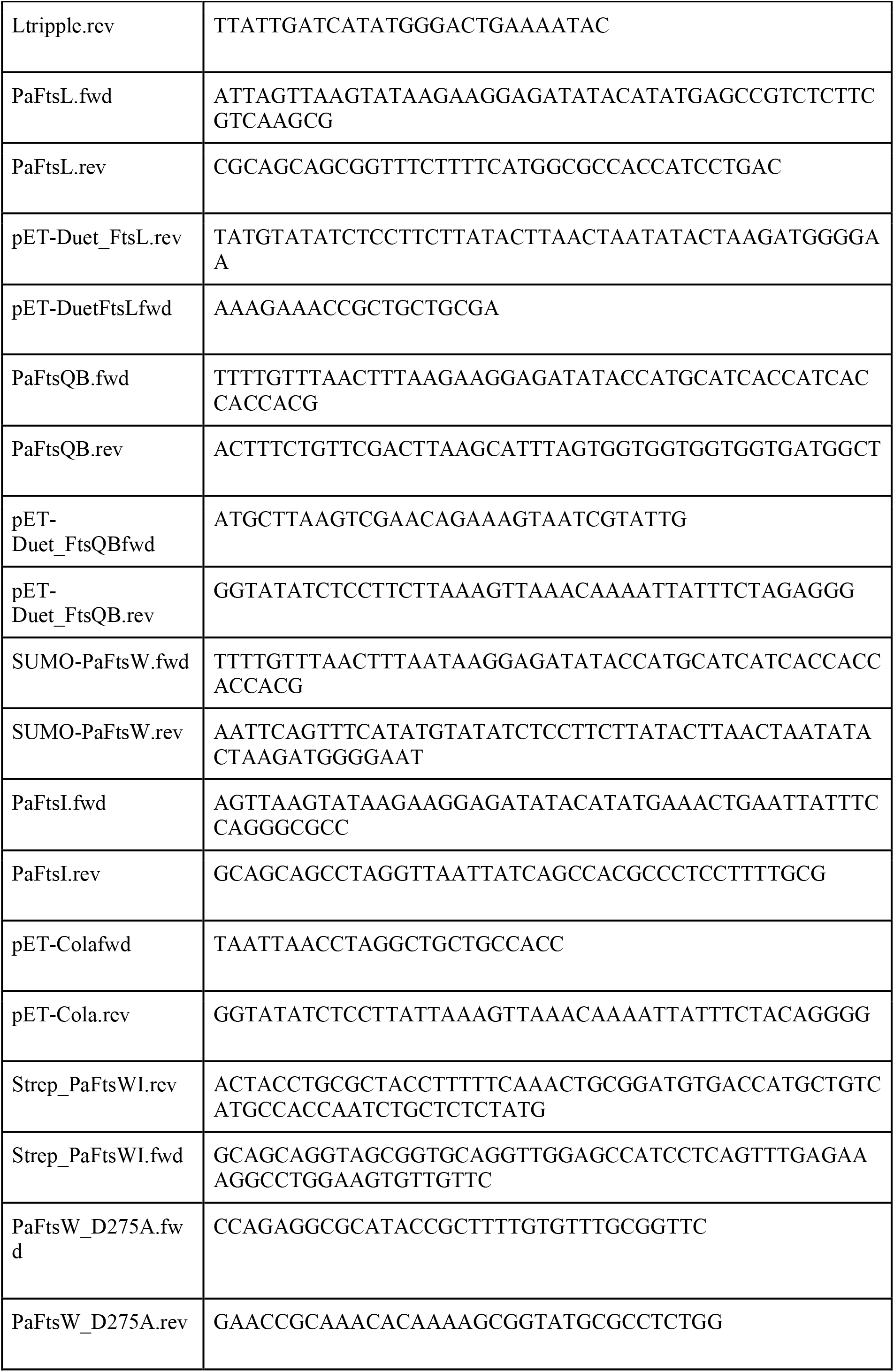
Primers.

**Table S2:**
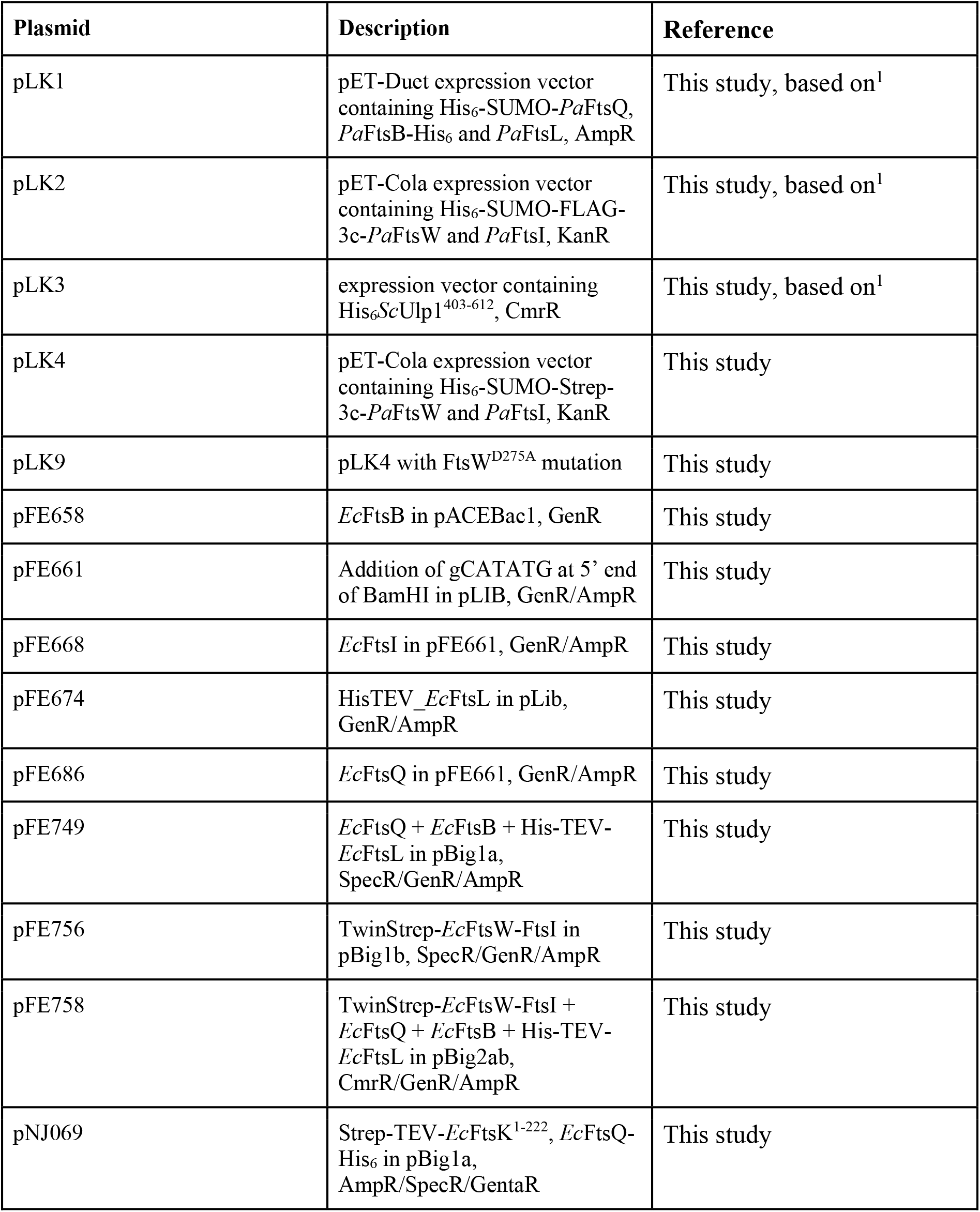
Plasmids.

| Plasmid | Description | Reference |
| --- | --- | --- |
| pLK1 | pET-Duet expression vector containing His<sub>6</sub>-SUMO-*PaFtsQ*, *PaFtsB*-His<sub>6</sub> and *PaFtsL*, AmpR | This study, based on<sup>1</sup> |
| pLK2 | pET-Cola expression vector containing His<sub>6</sub>-SUMO-FLAG-3c-*PaFtsW* and *PaFtsI*, KanR | This study, based on<sup>1</sup> |
| pLK3 | expression vector containing His<sub>6</sub>*ScUlp1*<sup>403-612</sup>, CmrR | This study, based on<sup>1</sup> |
| pLK4 | pET-Cola expression vector containing His<sub>6</sub>-SUMO-Strep-3c-*PaFtsW* and *PaFtsI*, KanR | This study |
| pLK9 | pLK4 with *FtsW*<sup>D275A</sup> mutation | This study |
| pFE658 | *EcFtsB* in pACEBac1, GenR | This study |
| pFE661 | Addition of gCATATG at 5' end of BamHI in pLIB, GenR/AmpR | This study |
| pFE668 | *EcFtsI* in pFE661, GenR/AmpR | This study |
| pFE674 | HisTEV-*EcFtsL* in pLib, GenR/AmpR | This study |
| pFE686 | *EcFtsQ* in pFE661, GenR/AmpR | This study |
| pFE749 | *EcFtsQ* + *EcFtsB* + His-TEV-*EcFtsL* in pBig1a, SpecR/GenR/AmpR | This study |
| pFE756 | TwinStrep-*EcFtsW*-*FtsI* in pBig1b, SpecR/GenR/AmpR | This study |
| pFE758 | TwinStrep-*EcFtsW*-*FtsI* + *EcFtsQ* + *EcFtsB* + His-TEV-*EcFtsL* in pBig2ab, CmrR/GenR/AmpR | This study |
| pNJ069 | Strep-TEV-*EcFtsK*<sup>1-222</sup>, *EcFtsQ*-His<sub>6</sub> in pBig1a, AmpR/SpecR/GentaR | This study |

**Figure S1:**
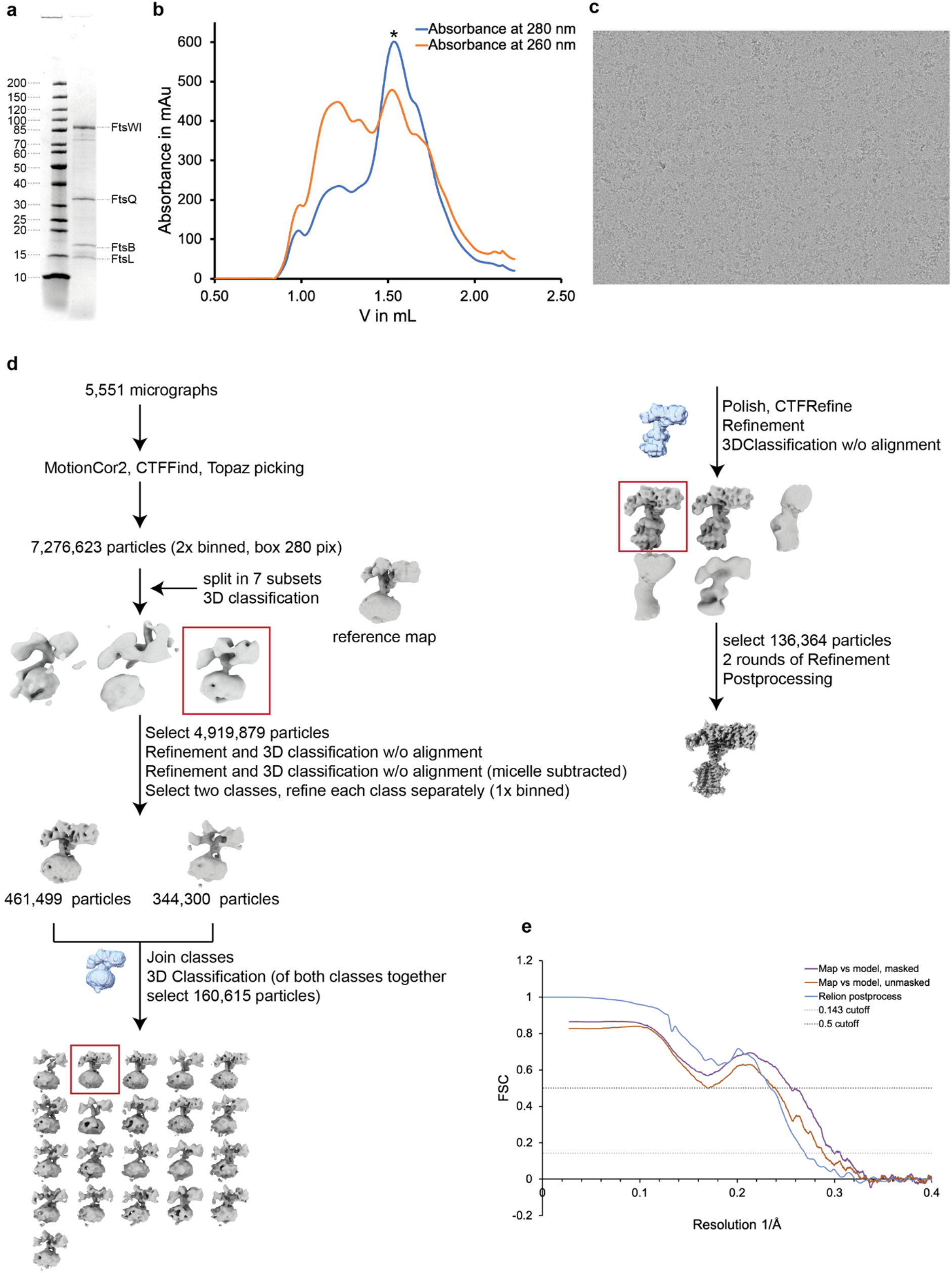
Protein purification and cryo-EM data processing. **a**) SDS-PAGE gel of the co-expressed and purified *Ec*FtsWIQBL divisome core complex after size-exclusion chromatography. FtsW and FtsI are joined by a short linker in this construct. **b**) Size-exclusion chromatogram of *Pa*FtsWIQBL. The measured absorbances at 260 nm and 280 nm are shown in orange and blue, respectively. The peak fraction of the size exclusion run, indicated with an asterisk, was used for grid preparation. **c**) Representative micrograph of *Pa*FtsWIQBL used for the final reconstruction. **d**) Cryo-EM processing scheme for *Pa*FtsWIQBL. **e**) Fourier Shell Correlation (FSC) curves for the *Pa*FtsWIQBL cryo-EM maps and structures.

**Figure S2:**
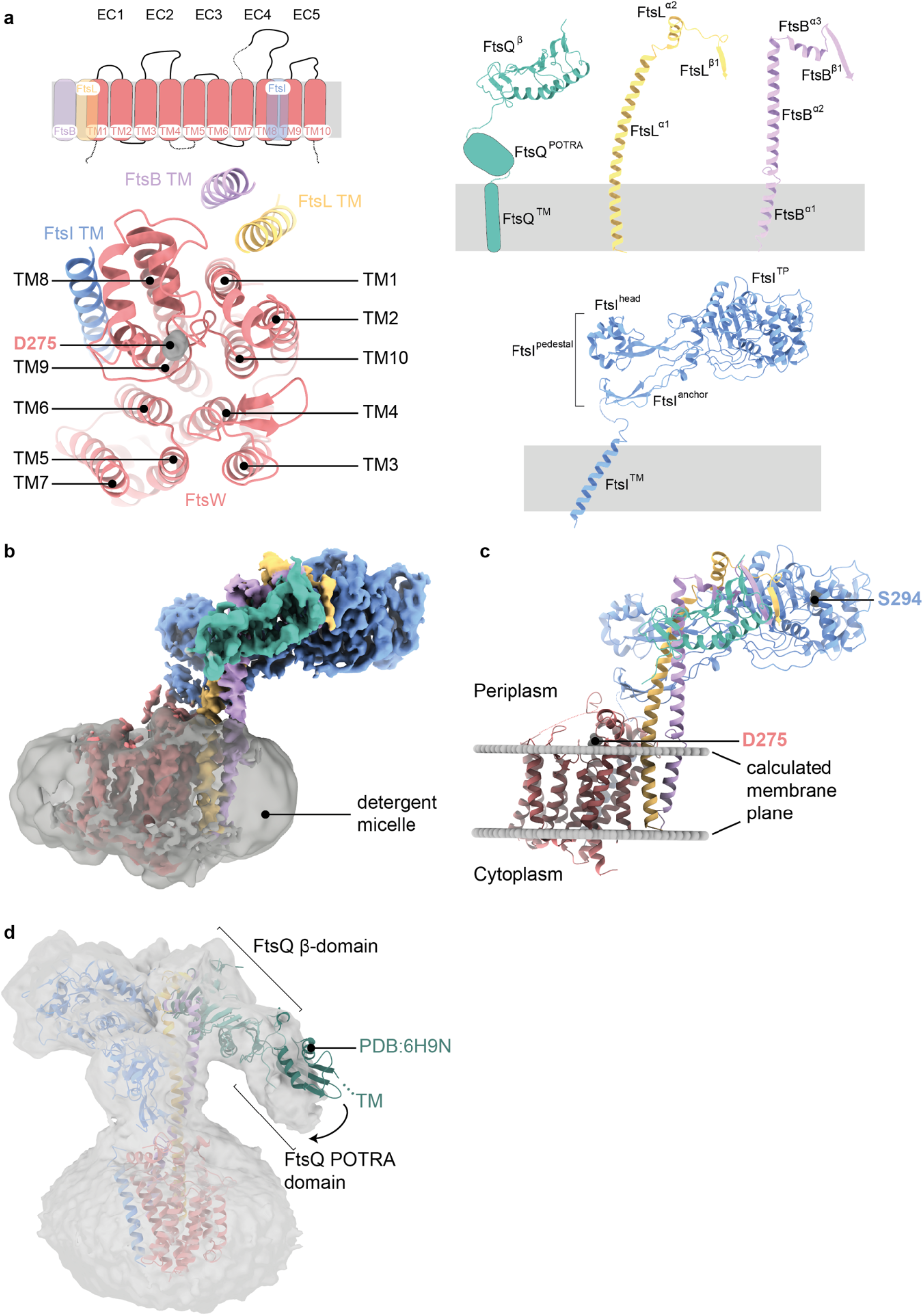
Architecture of the *Pa*FtsWIQBL complex. **a**) Upper left panel: schematic of the transmembrane helices of FtsW, FtsI and FtsL. Two extracellular loops of FtsW that could not be build due to missing density and the N- and C-terminal tails of FtsW are indicated by doted lines. Lower left panel: top view of the transmembrane domain, with FtsW transmembrane helices consecutively numbered based on the sequence (identical to numbering of helices in a previous RodA structure^21^). FtsW’s putative active site residue D275 is indicated. Right panel: Labelling of the different domains in FtsQ, FtsL, FtsB and FtsI that was used throughout the paper. **b**) Cryo-EM density showing *Pa*FtsWIQBL within the Lauryl Maltose Neopentyl Glycol (LMNG) detergent micelle, which was subtracted during the later processing stages. **c**) Prediction of the position and orientation of the divisome core complex transmembrane segments in the lipid bilayer using the Orientations of Proteins in Membranes webserver^27^. The membrane plane is indicated with two grey discs and the active sites of FtsW and FtsI are labelled. **d**) A low-resolution structure obtained after fewer 3D classifications shows additional density for FtsQ^POTRA^ at low contour levels and indicates that the transmembrane segment of FtsQ is most likely not part of the micelle that contains the other TM segments. Alignment of a previous FtsB:FtsQ crystal structure (PDB: 6H9N) on FtsQ^β^ shows that FtsQ^β^ and FtsQ^POTRA^ adopt different conformations relative to each other.

**Figure S3:**
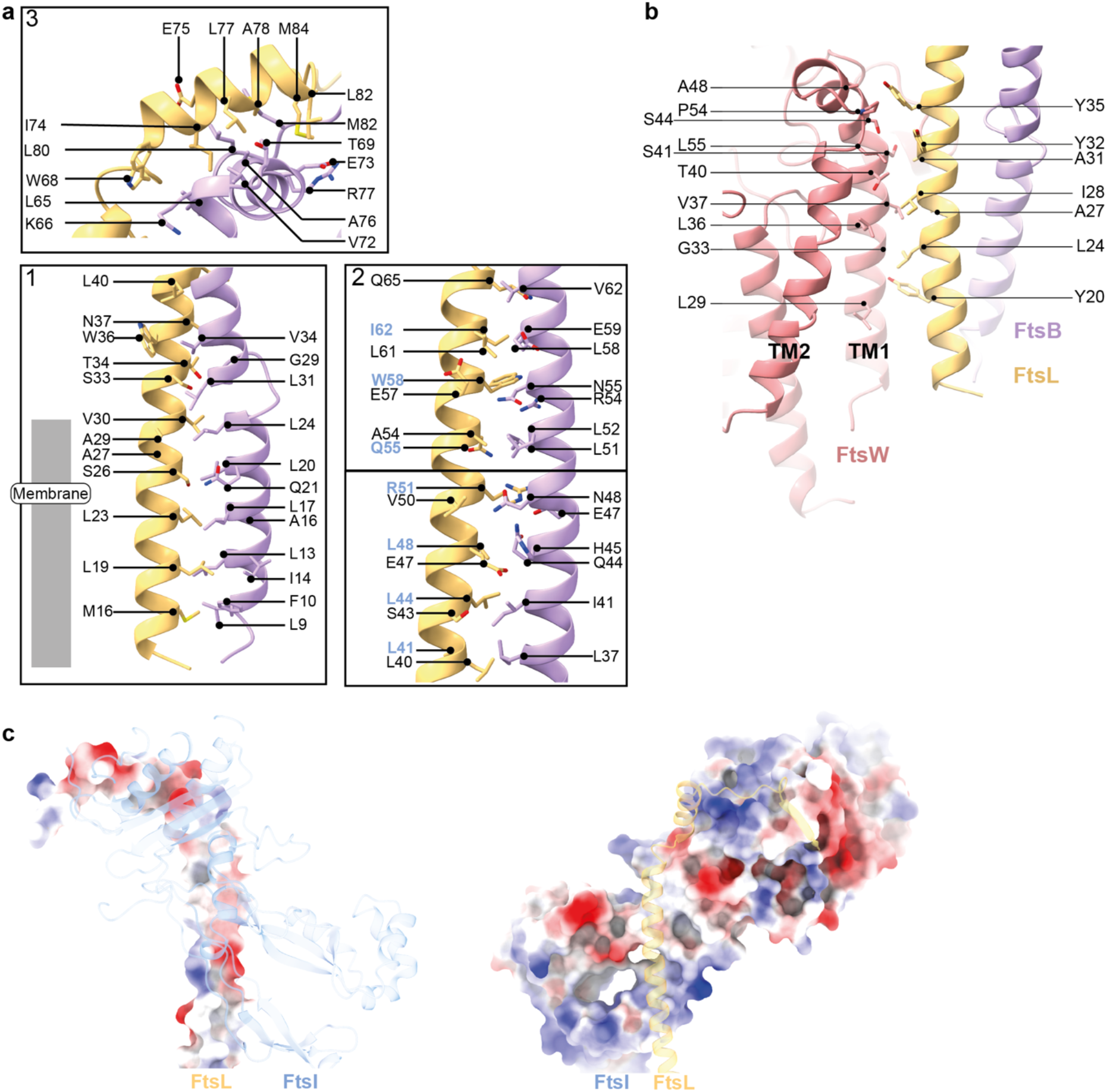
Detailed analysis of the interactions between FtsB and FtsL, FtsW and FtsL and FtsI and FtsL. **a**) Interaction sites between FtsB and FtsL, as also indicated in Figure 2a. Residues of FtsL that also interact with FtsI are highlighted in blue. **b**) Analysis of the interaction sites between FtsL and FtsW in their transmembrane region. The coiled coil conformation of FtsL means that the interaction surface is not as extended as it would be if it were straighter and not in a coiled coil. **c**) Electrostatic analysis of the interactions between FtsI and FtsL shows that the interaction site in the coiled coil area is mainly hydrophobic/neutral.

**Figure S4:**
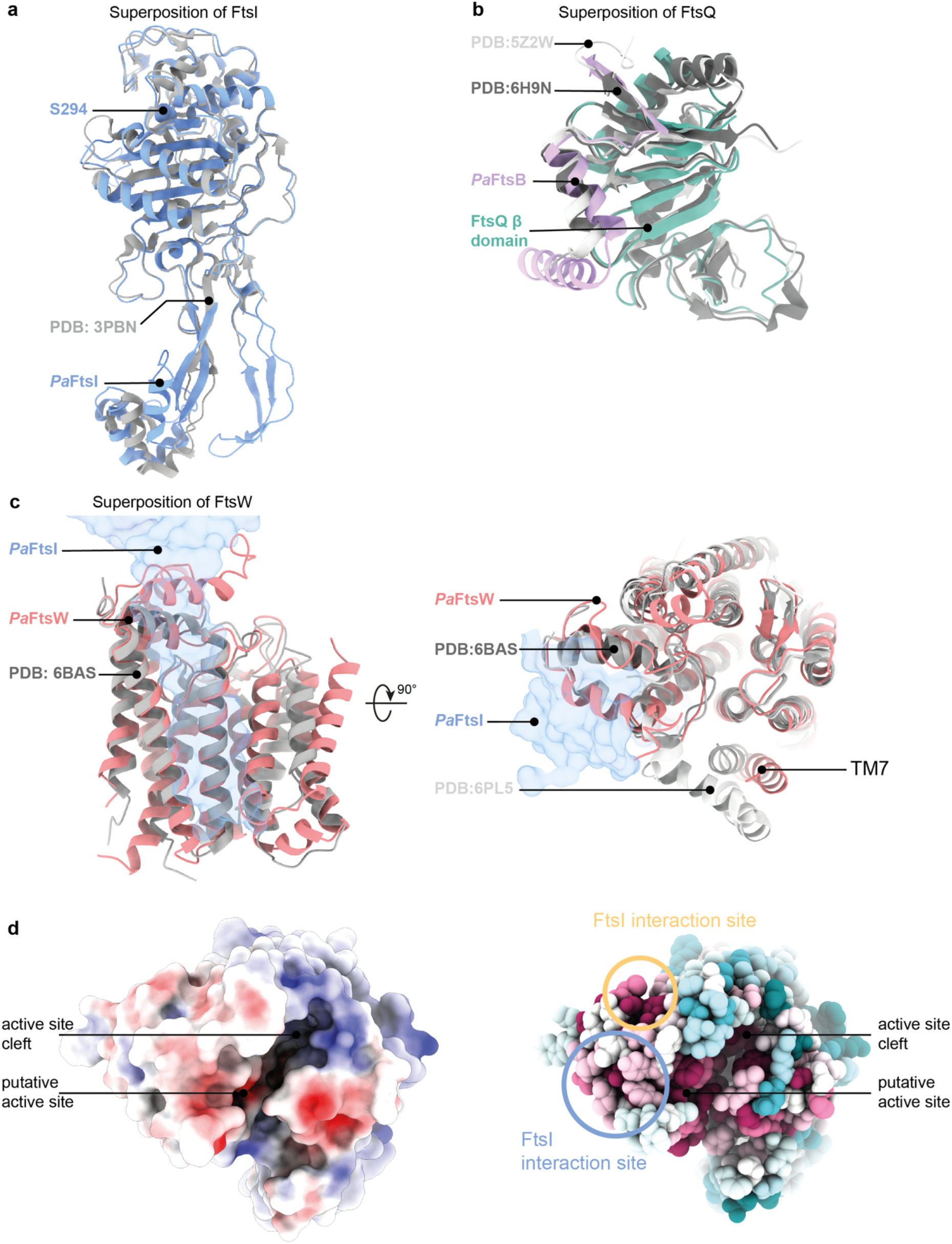
Comparison of *Pa*FtsWIQBL cryo-EM structure with previous crystal structures of FtsI, FtsQ and FtsW/RodA. **a**) Superposition of FtsI determined by X-ray crystallography (PDB: 3PBN, grey) with the FtsI part of PaFtsWIQBL cryo-EM structure. The TP active site residue S294 is indicated (RMSD of 0.708 Å across 372 pruned atom pairs). **b**) Superposition of FtsQB determined by X-ray crystallography (PDB: 6H9N in dark grey, PDB: 5Z2W in light grey) with the same area in the cryo-EM structure determined here (For alignment of FtsQ: RMSD (FtsQ-6H9N) of 1.186 Å across 86 pruned atom pairs, RMSD (FtsQ-5Z2W) of 1.118 Å across 95 pruned atom pairs). **c**) Superposition of RodA determined by X-ray crystallography (PDB: 6BAS in dark grey (left and right), PDB: 6PL5 in light gray (right)) and FtsW in the cryo-EM structure. The position of FtsI is indicated as a transparent blue outline. Apart from transmembrane helix 7, the structures align very well (RMSD (FtsW-6PL5) of 1.188 Å across 202 pruned atom pairs; RMSD (FtsW-6BAS) of 1.126 Å across 206 pruned atom pairs). **d**) Electrostatic surface representation of *Pa*FtsW viewed from the periplasmic side. A deep cleft is visible that contains the putative active site residue D275. The same representation showing sequence conservation of FtsW mapped onto the surface representation shows that this cleft is highly conserved. Additionally, interaction sites with FtsI and FtsL are indicated; these also show above average levels of sequence conservation.

**Figure S5:**
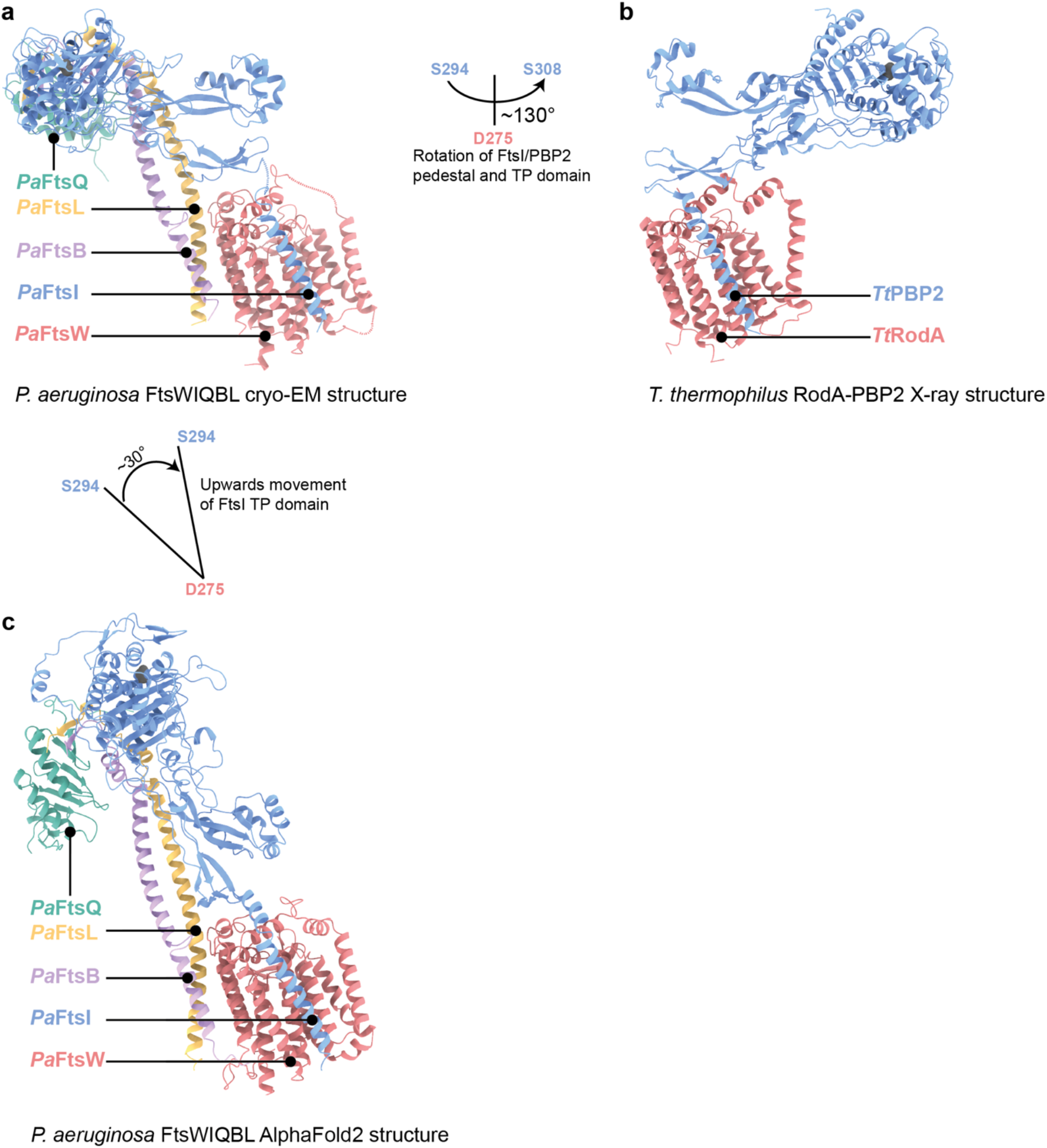
Comparison of the *Pa*FtsWIQBL cryo-EM structure with RodA-PBP2 structure and AlphaFold2 structure prediction. Comparison of the cryo-EM structure *Pa*FtsWIQBL **(a)**, the *Thermus thermophilus* RodA-PBP2 crystal structure (PDB: 6PL5, **b)** and the AlphaFold2 prediction of *Pa*FtsWIQBL **(c)**. All three structures were aligned on FtsW/RodA. The FtsQ^POTRA^ and FtsQ™ of the AlphaFold2 model were removed for clarity. The FtsI/PBP2 periplasmic domains show a large 130° lateral rotation between the *P. aeruginosa* FtsWI and *T. thermophilus* RodA-PBP2 models (a-b). The rotation was measured around an axis perpendicular to the membrane plane and intersecting the FtsW active site. The distance between both active sites in FtsI (S294) and PBP2 (S308) is 125 Å. A 30° vertical rotation of the periplasmic FtsI domains is visible between the cryo-EM and AlphaFold2 models of *Pa*FtsWIQBL (a-c). The angle was measured between the FtsW active site (D275) and the FtsI active sites (S294). The distance between the FtsI active sites in the cryo-EM and AlphaFold2 models is 46 Å.

**Figure S6:**
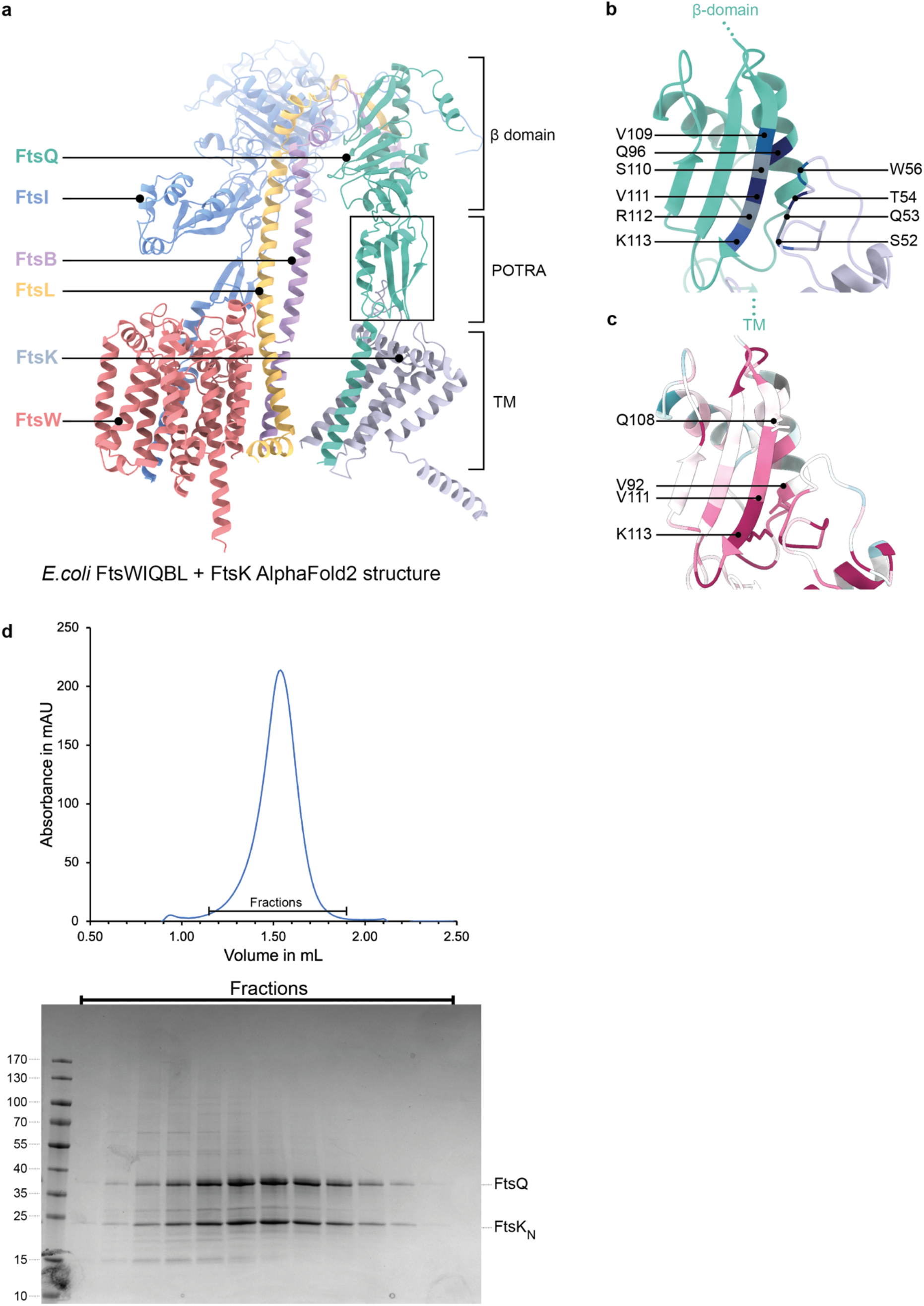
Interactions of FtsQ with FtsK. **a**) AlphaFold2 model of *E. coli* FtsWIQBL + FtsK N-terminal domain (residues 1-222; FtsK^1-222^, grey), showing a predicted interaction between FtsQ^POTRA^ and a periplasmic loop from FtsK (FtsK^W51-H57^). **b**) Co-evolutionary coupling analysis calculated with EVcouplings^28^ finds six out of the ten residue pairs located between FtsQ^POTRA^ and FtsK^W51-H57^ in the AF2 model in a): FtsQ^V109^ – FtsK^W56^ (blue), FtsQ^Q96^ – FtsK^T54^ (dark blue), FtsQ^S110^ – FtsK^Q53^ (gray), FtsQ^V111^ – FtsK^T54^ (dark blue), FtsQ^R112^ – FtsK^Q53^ (grey), and FtsQ^K113^-FtsK^S52^ (blue). **c**) Sequence conservation analysis (calculated using ConSurf webserver^29^) of the same area shows that the β-strands of FtsQ and FtsK^W51-H57^ that are predicted to interact are highly conserved. Amino acid residues that abolish FtsQ localisation (which is dependent on FtsK septum localisation in cells) when mutated are shown as sticks and are labelled^30^. **d**) Size-exclusion trace and SDS-PAGE gel of the co-expression and purification of *E. coli* FtsQ and FtsK^1-222^ shows clear co-migration of the proteins.

**Table S1:** Imaging statistics cryo-EM.

| **Data collection and processing** | ***PaFtsWIQBLEMD-16042*, PDB 8BH1** |
| --- | --- |
| Magnification | 75,000x |
| Voltage (kV) | 300 |
| Electron fluency (e-/Å<sup>2</sup>) | 41 |
| Defocus range (μm) | -1.2 to -3 |
| Pixel size (Å) | Nominally 1.09, refined to 1.05 |
| Symmetry | C1 |
| Initial particles (number) | 7,276,623 |
| Final particles (number) | 136,364 |
| Map resolution (Å) | 3.7 |
| FSC threshold | 0.143 |
| **Model** |  |
| Initial model used | AlphaFold2 model |
| Model resolution (Å) | 3.9 |
| FSC threshold | 0.5 |
| Map sharpening B factor (Å<sup>2</sup>) | -127 |
| *Model composition* |  |
| Non-hydrogen atoms | 9,269 |
| Protein residues | 1,195 |
| RMSD bond lengths (Å) | 0.004 |
| RMSD bond angles (°) | 0.722 |
| *Validation* |  |
| MolProbity score | 1.98 |
| Clashscore | 16.71 |
| Rotamers outliers (%) | 0.0 |
| Ramachandran plot |  |
| Favored (%) | 96.18 |
| Allowed (%) | 3.82 |
| Disallowed (%) | 0 |

